# PIPENN: Protein Interface Prediction with an Ensemble of Neural Nets

**DOI:** 10.1101/2021.09.03.458832

**Authors:** Bas Stringer, Hans de Ferrante, Sanne Abeln, Jaap Heringa, K. Anton Feenstra, Reza Haydarlou

## Abstract

**Motivation:** Protein interactions play an essential role in many biological and cellular processes, such as protein–protein interaction (PPI) in signaling pathways, binding to DNA in transcription, and binding to small molecules in receptor activation or enzymatic activity. Experimental identification of protein binding interface residues is a time-consuming, costly, and challenging task. Several machine learning and other computational approaches exist which predict such interface residues. Here we explore if Deep Learning (DL) can be used effectively for this prediction task, and which learning strategies and architectures may be most efficient. We introduce seven DL architectures that are applied to eleven independent test sets, focused on the residues involved in PPI interfaces and in binding RNA/DNA and small molecule ligands.

**Results:** We constructed a large data set dubbed `BioDL`, comprising protein-protein interaction data from the PDB and protein-ligand interactions (DNA, RNA and small molecules) from the BioLip database. Additionally, we reused our existing curated homo- and heteromeric PPI data sets. We performed several experiments to assess the impact of different data features, spatial forms, encoding schemes, network initializations, loss functions, regularization mechanisms, and activation functions on the performance of the predictors. Benchmarking the resulting DL models with an independent test set (`ZK448`) shows no single DL architecture performs best on all instances, but that an ensemble of DL architectures consistently achieves peak prediction performance. Our PIPENN’s ensemble predictor outperforms current state-of-the-art sequence-based protein interface predictors on all interaction types, achieving AUCs of 0.718 (protein–protein), 0.823 (protein–nucleotide) and 0.842 (protein– small molecule) respectively.

**Availability:** Source code and data sets at https://github.com/ibivu/pipenn/

## 1 Introduction

Protein interactions are crucial in many biological and cellular processes [1], such as transcription, signal transduction or enzymatic activity. Proteins, through their binding interfaces, interact with each other and a variety of other molecules, giving rise to all manner of cell functions. Knowledge about these interfaces provides essential clues about the mechanisms underlying associated activities. This molecular level knowledge can be obtained by experimental and computational methods, and is applied in many scientific and therapeutic areas [2].

Protein interaction prediction refers to a set of computational methods that aim to predict protein bindings of different interaction types (protein– protein (PPI), protein–small molecule, protein–nucleotide (DNA/RNA) (see SI Fig. S1) at different binding levels (binding-site, binding-residue) [3, 4, 5, 6]. In general, such methods are based on information from protein structure and/or sequence, and may utilize various classic Machine Learning (ML) [7, 8, 9, 10] and Deep Learning (DL) architectures [11, 12]. DL architectures are composed of multiple building blocks, such as initialization, regularization, loss, activation, each containing various parameters. To predict protein bindings of the mentioned interaction types at residue-level, using only information related to protein sequence, we explore the following question: Which DL architecture and composition of architectural building blocks is able to improve the performance of predictors? Hence, we designed several DL models based on the following types of neural net architectures.

The simplest neural architecture considered here is the fully connected *Artificial Neural Network* (ANN): every neuron in a layer is connected to all neurons in the next layer. This configuration is general-purpose and structure agnostic. The input to this network is a single protein residue at a time, which means their sequence context is not considered.

*Convolutional Neural Network* (CNN) architectures are already extensively used in various flavours in computational biology e.g. [11], including protein interaction site prediction e.g. [4]. A CNN consists of three types of hidden layers: *convolutional layers*, *pooling layers*, and *fully connected layers*. For predictions in a protein sequence, neurons are organized sequentially (1D spatial form) and every neuron is connected to the neurons of a local region (receptive field) in the previous layer. Pooling layers perform down-sampling to speed up computation, and fully-connected layers perform the actual classification task based on the abstract representations of the original protein sequence input.

*Dilated Convolutional Networks* (DCN) achieve large receptive fields by gradually increasing the dilation rate in subsequent layers [13]. This allows DCNs to remain relatively shallow, require few parameters and converge quickly. DCNs typically maintain high output resolutions without up-sampling, and have been successfully applied in many areas [14], including computational genomics [15, 16].

The *U-Net* is the most commonly used CNN architecture, especially when the the amount of training data is limited [17]. In a U-Net, high-dimensional information is first reduced (contracted) to a smaller latent space, and subsequently increased (expanded) to the original dimensionality. Visually, this gives rise to a U-shape, from which this architecture derives its name. Though more typically used in image recognition tasks, we find U-Nets – like CNNs in general – equally applicable for 1D (sequence-based) predictions.

In general, deep CNNs can learn more complex features because of having more layers in their architectures. However, it does not mean that deeper networks will always perform better than their shallow counterparts. There is a maximum threshold for the number of layers and this maximum can be easily reached, specially, in traditional CNNs. The most important reason for this limitation is the problem of vanishing or exploding gradient, where the network weights of the first layers can no longer be correctly updated. As a result, the network is not anymore able to propagate the information to the higher layers. To cope with this, [18] proposed a residual learning framework (*ResNet*).

*Recurrent Neural Networks* (RNN) are particularly suitable for learning non-linear dependencies in sequential data, such as protein sequences. No limit is imposed on the size of the input series of amino acid symbols, but each symbol is represented by additional layers. To mitigate the vanishing/exploding gradient in the resulting many-layer models [19] and retain a computationally efficient training, *Gated Recurrent Unit* (GRU) was proposed by [20]. [21] showed that the performance of GRU is on par with the more complex LSTM [19] on sequence modeling tasks, while taking much less time to train.

Recently, hybrid architectures of CNNs and RNNs are increasingly employed in computational biology. The hybrid models aim to get the best of both worlds: the spatial aspects of CNNs combined with the temporal aspects of RNNs. For instance, [12] combined residual convolutional network with LSTM to predict a protein’s residue-residue contacts (contact map prediction task). They use *ResNet* to capture spatial relationships between local residues, and LSTM to capture long-range relations between non-local residues. In the field of genomics, [22] integrated CNNs with bi-directional LSTMs for modeling of the properties and functions of non-coding DNA sequences. To learn a regulatory grammar in the DNA motifs, they use convolution layers to capture local patterns in the motif sequences, and recurrent layers to capture long-term dependencies between the motifs.

Here, we explore the effectiveness of each of the above neural net architectures: *ann*, *dnet*, *unet*, *rnet*, *rnn*, and *cnet*. We expect they will each capture overlapping but distinct patterns in protein sequence data, yielding different predictions for which residues are part of an interface. Finally, we include an ensemble architecture, which combines the outputs of these six neural nets into one. All models, collectively referred to as PIPENN, are trained on five different training sets. Their performance is benchmarked with 11 test sets, including the standardized benchmark data set introduced by [3], referred to as `ZK448`. The latter is also used to assess and compare the performance of the PIPENN ensemble method with competing models. To our knowledge, we are the first to apply DL architectures to different types of protein interface prediction at this scale, and obtain the best prediction results to date.

## 2 Materials and methods

### 2.1 Data sets

To perform experiments on different types of protein interaction data, i.e. PPI, small molecule, and nucleotide (DNA/RNA), we used five data sets for training (see SI Table S1) and 11 independent data sets for testing (see SI Table S2). The `HHC` data set, containing homo- and heteromeric proteins annotated with PPI interface residues, was already available in our group from [8, 9]. Another part was obtained from [3]: the benchmark test set `ZK448`, which contains proteins annotated with residues for all types of protein interaction. The final part was newly constructed: the BioDL data set that contains proteins annotated with residues for all types of protein interaction, as described below and illustrated in Figure 1.

**Figure 1:**
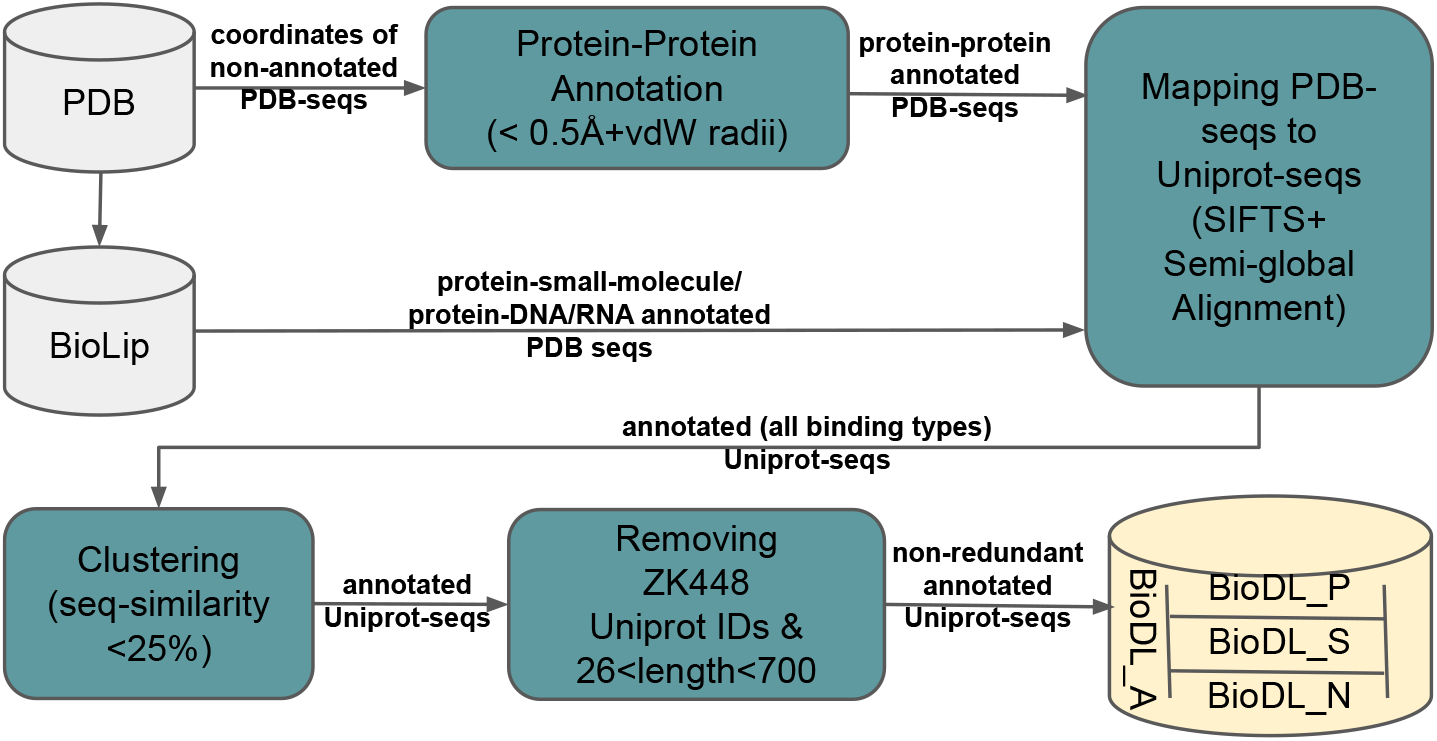
Generation of the `BioDL` data set from the PDB and BioLip databases.

For small molecule and nucleotide interactions, we retrieved the whole BioLip [23] database and extracted the interaction annotation data. For PPIs, we downloaded the coordinates of 138 729 protein structures of 2.5 Å resolution or lower, excluding fragments, from the PDB [24] on 3 April 2019. Following the annotation criterion in BioLip, we annotated a residue as interacting if the distance between one of its atoms and any atom of the ligand is less than the sum of their Van der Waals radii plus 0.5 Å.

The newly derived annotations and the existing annotations from BioLip are associated with PDB sequences. In order to test our models on the benchmark test set `ZK448`, where annotations are associated with Uniprot sequences, we mapped PDB sequences to Uniprot as follows. We used SIFTS [25] to retrieve all Uniprot sequences corresponding to the PDB entries. Entries with missing or conflicting Uniprot IDs were discarded. We mapped residues between PDB and Uniprot sequences by alignment with harsh penalties (mismatches −5, gap opening −20 and extension −50). Alignments where less than 80% of interaction site residues were mapped, or with more than two inserts, five deletions, three mismatches in the interaction sites, or five mismatches were discarded.

Subsequently, `BLASTClust` was used to cluster the obtained Uniprot sequences at 25% sequence similarity. All clusters containing a Uniprot ID used in `ZK448` were removed and thereafter randomly one sequence from each cluster was chosen. From this non-redundant data set, based on the criterion used by [26], proteins having sequences longer than 700 and shorter than 26 amino acids were removed. The final data set was split into a training set `BioDL_A_TR` (95%) and a testing set `BioDL_A_-TE` (5%), containing annotations for all three types of interactions. We further split these into proteins annotated with PPI (`BioDL_P_TR &_TE`), small molecule (`BioDL_S_TR &_TE`), and nucleotide (`BioDL_N_TR &_TE`) interactions. See SI Table S1 and S2 for data set statistics of the training (`TR`) and test (`TE`) sets, respectively.

### 2.2 Data features

Our models predict various types of interactions for a given single protein sequence. For each residue along the sequences in our BioDL data sets, we use the global feature protein length and the following local features: amino acid type (AA), Position Specific Scoring Matrix (PSSM), accessible surface area (SA), secondary structure (SS), and conserved domain.

Protein length is the number of residues in the Uniprot sequence of the query protein. `PSI-BLAST` [27] was utilized to obtain PSSM profiles for each sequence using max. 500 sequence homologs from the NR70 database, using three iterations and an E-value threshold of 0.001. As a result, we obtained 20 PSSM values (one per amino acid type) for each residue. The PSSM scores were normalized using the sigmoid function. Further, we used `NetSurfP` to predict from sequence the Absolute/Relative Surface Accessibility (ASA/RSA) and the propensity for *α*-helix (PA), *β*-sheet (PB), and coil (PC) [28]. To indicate whether or not a residue belongs to a conserved protein domain, we employed the protein domain information from the Pfam [29] database for the Uniprot sequences.

Finally, we included four aggregated features for each of the features. Each aggregated feature is the average of the feature value over a specific number of the neighbouring residues: 3, 5, 7, or 9. They are abbreviated as `<winsize>_wm_<fname>`, e.g., `9_wm_PB` refers to the average of the predicted *β* sheet probabilities over a window of size 9. This leads to a total of 128 features, as detailed in SI Table S13.

### 2.3 Learning architectures

The input layer of our ANN based architecture *ann* consists of a number of neurons, each representing a feature of one amino acid (see SI Fig. S2).

The network has eight hidden layers and its output is a binary classification predicting whether or not an amino acid is part of an interface. In order to capture as much as possible information about a residue, the number of neurons in the first hidden layer is the highest, decreasing gradually in the subsequent layers to achieve more general representations. However, it has a large number of parameters (weights) and relatively long convergence time (see SI Table S11).

We designed three different CNN architectures *dnet*, *unet*, and *rnet*. The input layers for these consist of one neuron per feature per amino acid in the protein sequence. *dnet* is based on dilated CNN, has five *convolutional layers*, zero *pooling layer*, and one *fully connected layer* (SI Fig. S3). The dilation rate increases from one to 16. *unet* is based on the U-Net architecture and the contraction block starts with 1024 (length of padded protein sequence) and is gradually down-sampled by max-pooling to 32 at the bottleneck (SI Fig. S4). For up-sampling *transposed convolutions* are used [30]. *rnet* is a residual CNN that allows us to experiment with deeper CNNs (SI Fig. S5). We use a slightly modified version of the *full pre-activation* configuration [31]. In *rnet*, a residual block consists of two sequential *full pre-activation* configurations, each containing a *Dropout*, *Batch-Normalization*, and *Parametric Rectified Linear Unit* (PReLU) followed by a 1D convolution. We use eight such residual blocks.

Our recurrent NN architecture *rnn* consists of two GRU layers, each containing 1024 cells with each cell having an output dimension of 128 that goes to the next cell (SI Fig. S6). Finally, our *cnet* (SI Fig. S7) is a hybrid architecture that simply combines our *rnet* residual and *rnn* recurrent architectures.

### 2.4 Architecture building blocks and parameters

DL architectures are composed of multiple building blocks, each containing various parameters. Different compositions enable learning architectures to provide great flexibility and a broad application area. The choice for building blocks and parameter values strongly influences the performance that may be achieved, however, this depends both on the architecture and on the data set, making this one of the main challenges in DL. Below, we briefly motivate each of our choices.

- **Initialization method (IM):** We experimented with uniform and normal variants of the *Glorot* [32] and *He* [33] IMs. *He-Uniform* initialization proved most suitable for our architectures.
- **Regularization method (RM):** To overcome overfitting, we explored combinations of RMs: Lasso (L1), Ridge (L2), Batch-Normalization, Dropout, and Early-Stopping. A combination of *Batch-Normalization*, *Dropout* (20%), and *Early-Stopping* (based on the AUC metric) yielded best results.
- **Loss function (LF):** We explored a number of LFs: *Mean Squared Error* (MSE), *Jaccard*, *Tversky*, and *Cross-Entropy*. After running several experiments and due to having significant class imbalance in our data, we decided to use the *Cross-Entropy* loss with an additional term that compensates for class imbalance.
- **Activation function (AF):** Our choice of AF is based on a neuron’s position in the network, the computational speed of calculating its gradient, and its differentiability. We used *Sigmoid* at the last layer of all our architectures. For the hidden layers, using *Tanh* for the *rnn* architecture and *PReLU* for all other architectures provided best performance.
- **Encoding scheme (ES):** We performed experiments with *One-Hot* and *ProtVec* to encode amino acid sequences. *One-Hot* represents each amino acid simply as a 20-dimensional sparse vector. *ProtVec* constructs 3-gram words of amino acids for each protein sequence from *Swiss-Prot*, and trains a skip-gram neural network on this data [34]. The output of the network is an embedding space of 100-dimensional dense vectors, which we used for as input features for our models. We achieved better performance with the *One-Hot* representation.

### 2.5 Training and testing procedures

We split each of the training data sets into two parts: *train-set* and *val-set* (SI Table S1). The *train-set* part (80%) is used exclusively for training, and the *val-set* part (20%) is used for three purposes: (1) for gaining insight into the performance of an architecture, (2) for terminating the training when the performance does not show any improvement, and (3) for training of the *ensnet* architecture. The training procedure is as follows. After all six architectures are trained on a *train-set*, their trained models are applied on the corresponding *val-set*. The obtained predictions of each model become the training data for the *ensnet* architecture, see overview in Figure 2.

**Figure 2:**
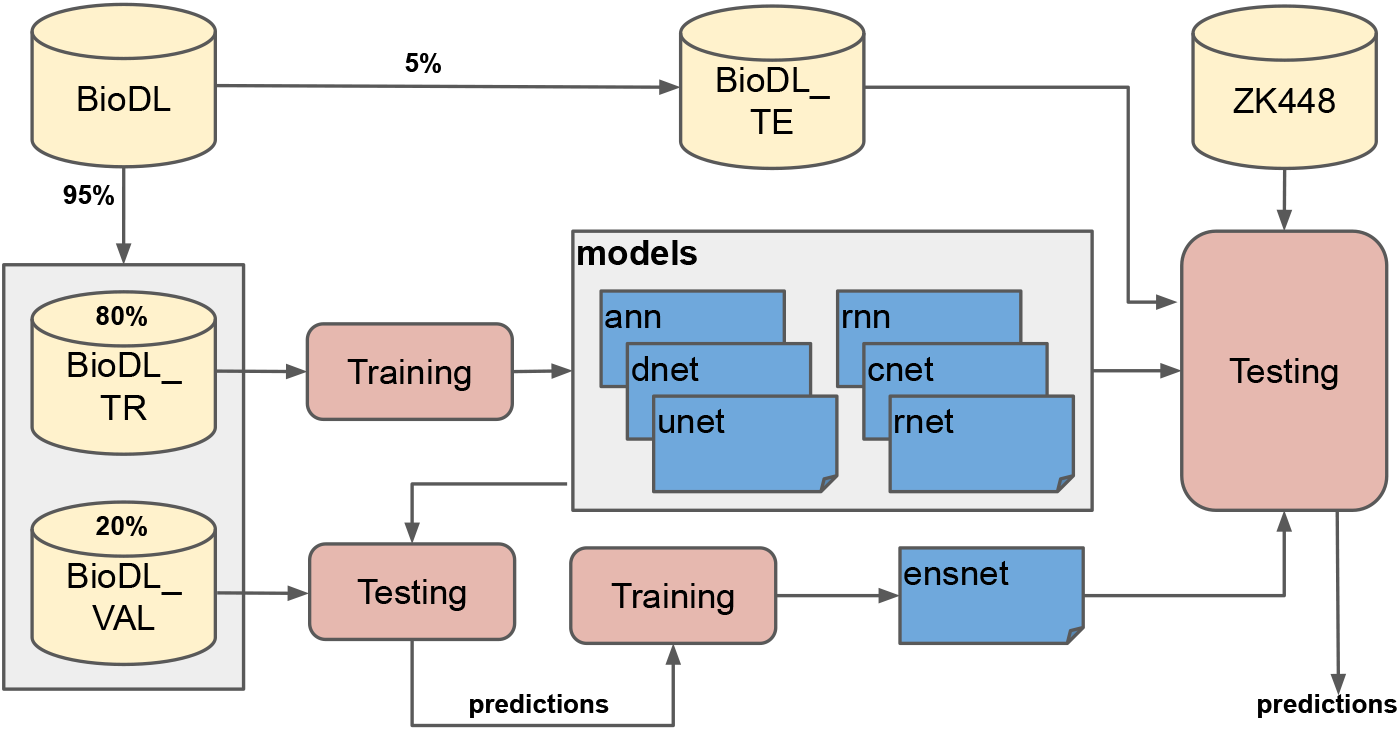
Training and testing procedure of our predictors (see Section 2.5 for the explanation of the procedure).

For all our architectures and data sets we used the same hyper-parameters, software platform, and infrastructure. The hyper-parameters that influence the performance and convergence time of the architectures are: batch size (8), optimization algorithm (*Adam*), learning rate (1e–4), maximum number of epochs (300), padding constants, bias vector usage, and the float size (64). We used the `Keras API of Tensorflow-2.1`.0 to build our architectures and trained them on a Linux machine having 32 CPUs, 2 GPUs, and 256 GB memory (see SI Table S11 for the run-time statistics).

We evaluated the performance of our trained models on the completely independent test sets shown in SI Table S2, using *Accuracy* (ACC), *Specificity* (SPEC), *Precision* (PREC), *Sensitivity* /*Recall* (SENS), *Balanced F-score* (F1), *Matthews Correlation Coefficient* (MCC), average precision on the PR-curve (AP), and area under the ROC curve (AUC). We calculate p-values for differences in AUC-ROC using the approach by [35]. Except for AP and AUCs, all metrics use the confusion matrix. MCC and AP are insensitive to class imbalance (on average about 11% of residues are interface, see SI Table S1). All results shown are based on the `Equal` method [3], by which a cutoff point is selected where FP and FN are equal. Hence, the values of PREC, SENS, and F1 are always the same. Output (predictions) of the models are unpadded before applying the metrics. Performance plots were created using the Plotly package.

### 2.6 Feature importance

For scoring and ranking the importance of features, we used the `KernelExplainer` from the SHAP package [36]. For each sample in the test set the contribution of each feature to the predicted outcome of a model is estimated, which is called a SHAP value.

## 3 Results

### 3.1 BioDL is sufficiently large for deep learning

DL architectures learn best when trained on a large data set. We constructed the `BioDL` data set based on BioLip and PDB, for training and testing of our various architectures. Within `BioDL` we have collected annotations of three specific types of protein interaction: PPI in `BioDL_P`, Small molecule ligands in `BioDL_S`, and Nucleic acid interaction in `BioDL_N. A` fourth set annotates any of these three interactions: `BioDL_A`. For better comparison with our previous work using Random Forest models, we also include a fifth data set: `HHC` from [9, 37]. Each of these data sets are further split into training (`_TR`) and test (`_TE`) sets as shown in Figure 2; see SI Table S1 for an overview and statistics of training sets. For `HHC_TE` we also report separately for homomeric (`Homo_TE`) and heteromeric (`Hetero_TE`) `PPI`. As further independent test sets we use `ZK448` from [3], which, like `BioDL`, contains protein, small molecule, and nucleotide interaction annotations (`ZK448_P_TE, ZK448_S_TE`, and `ZK448_N_TE`) as well as all combined (`ZK448_A_TE`). See SI Table S2 for an overview and statistics of test sets.

The overlap between the `BioDL` training sets is shown in a Venn di-agram in SI Fig. S13; out of 6 832 proteins in the training set there are only 85 with annotations for all three interaction types. This pattern is similar in the `BioDL` and `ZK448` test sets (SI Fig. S14).

We make distinctions between the models that were trained on the `BioDL_A_TR` data set containing all interaction types, i.e. *generic*, and those trained on a *type specific* interaction data, i.e. `BioDL_P_TR` for `PPI`, `BioDL_S_TR` for small molecules, and `BioDL_N_TR` for nucleic acids. We suffix those models correspondingly: −*a*, −*p*, −*s*, and −*n*. Models trained on the PPI-specific `HHC` data set are suffixed with −*hhc*.

### 3.2 Building block composition matters

We tested our predictors with different compositions of the architectural building blocks introduced in Section 2.4. These were trained on the smaller `HHC` PPI dataset for efficiency. Table 1 shows the metrics for different choices of building blocks for the *dnet hhc* predictor. Other architectures and data sets show very similar trends (data not shown). *dnet hhc* (the first row) corresponds to the composition with the highest performance in AUC: *HeUniform* kernel initialization, *CrossEntropy* loss function, *1D* spatial form, *PReLU* activation function, *Padding* for unifying the input shape, *One-Hot* amino acid encoding, and *Dropout* and *BatchNormalization* for regularization. Subsequent rows show the effect of substituting (→), adding (+) or removing (−) blocks. *GlorotNormal* kernel initialization yields highest average precision (AP); *MeanSquaredError* loss function highest accuracy (ACC), F1 and MCC. It is worth mentioning that for the best composition in *dnet hcc* we did not use the *MaxPooling*. All other variations yield lower performances, and the impact of omitting *Dropout*, *Padding*, and *BatchNormalization* is the strongest.

**Table 1:**
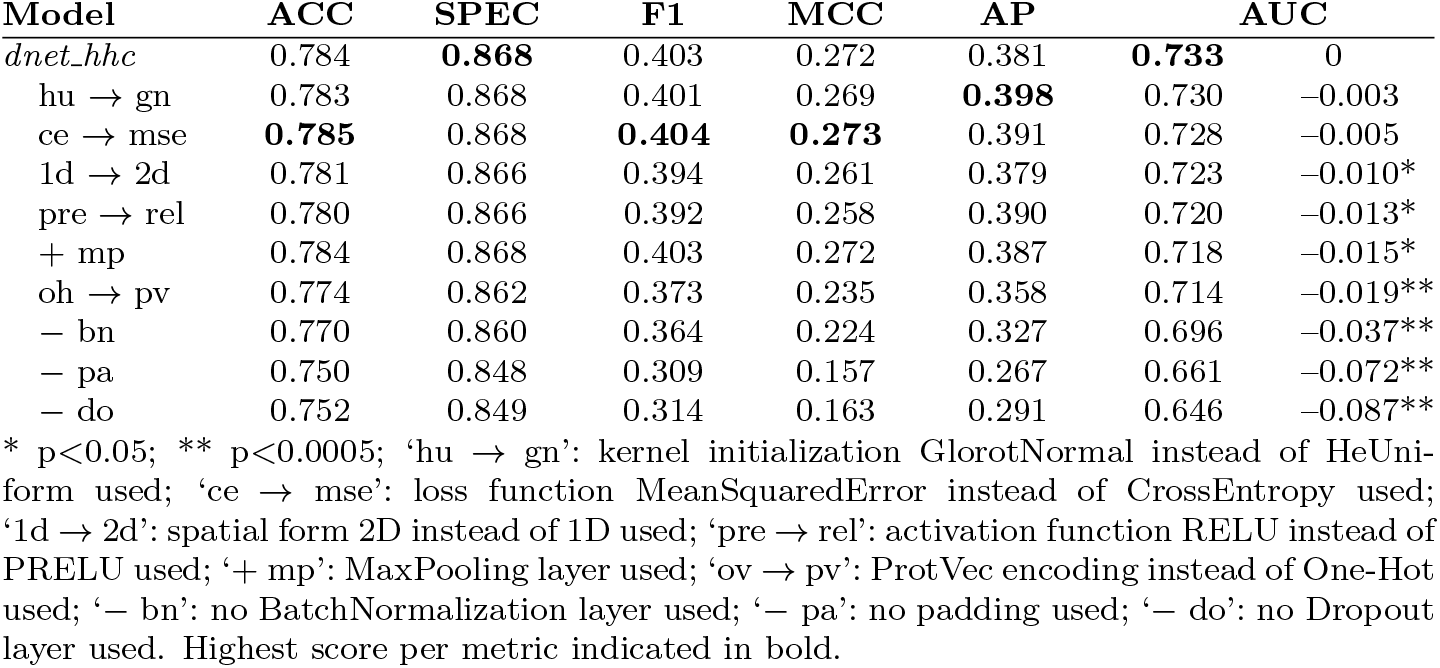
Impact of different architectural building blocks on the performance of the *dnet hhc* PPI predictor trained on `HHC_TR` and tested on `HHC_TE`.

### 3.3 Ensembling improves performance

We compared the performance of our ensemble predictor with those of our other DL predictors. Table 2 shows that *ensnet p* improves the AUC by 0.016 (p<0.0006) w.r.t. the best scoring other DL predic-tor (*dnet p*) on the PPI data, and it achieves the highest sensitivity (TPR) for any error (FPR) in the ROC plot (Figure 3A). Moreover, the P/R plot in Figure 3B shows that *rnet p* obtains the highest precision at low recall values but *ensnet p* achieves the highest precision at high recall values. To further investigate the reason behind this improvement, we explored the relation between accuracy of the *ensnet p* and the other predictors in scatter plots of MCC scores of *ensnet p* vs. the average and standard deviation of MCC scores of our six other predictors. Our analyses suggest that *ensnet* especially improves predictions for individual proteins where the *average* performance of the other models was already relatively high (SI Fig. S10).

**Table 2:**
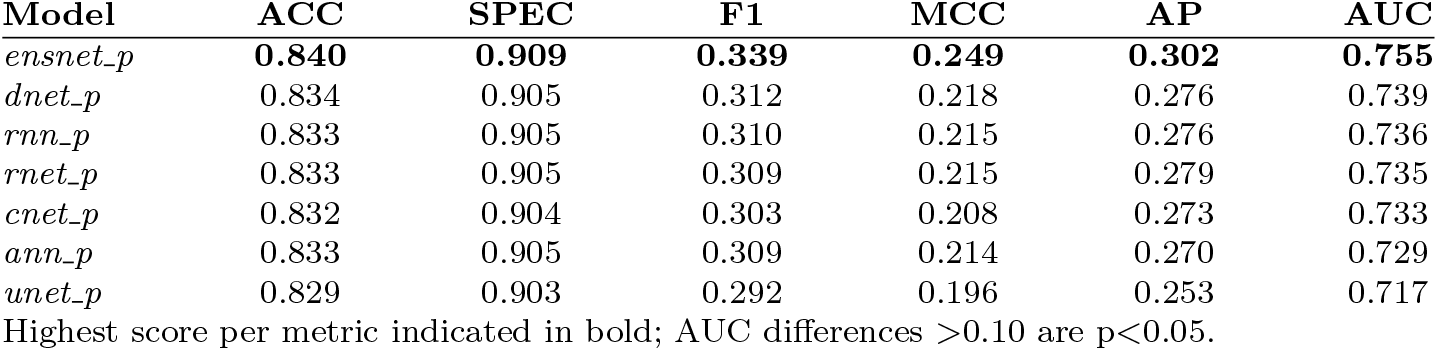
Performance of the ensemble PPI predictor *ensnet p* compared with all other predictors trained on `BioDL_P_TR`, and tested on `BioDL_P_TE`.

**Figure 3:**
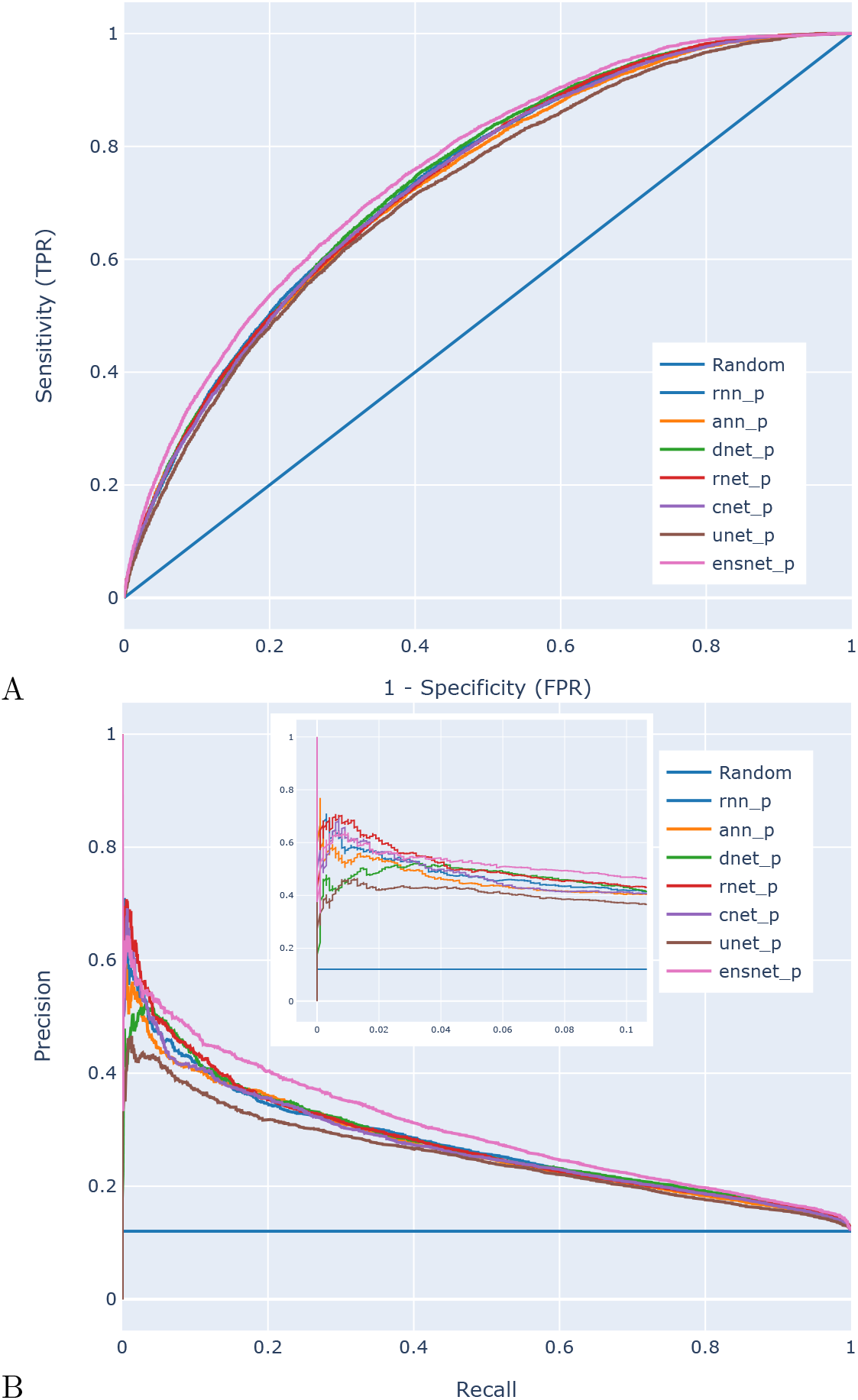
A ROC and B P/R plots of all six architecture models and the ensemble models, trained on `BioDL_P_TR` and tested on `BioDL_P_TE_PPI` data. The *ensnet p* clearly outperforms the six architecture models in the ROC plot, and in the P/R plot only *rnet p* and *rnn p* yield somewhat higher precision (~0.6) at very low recall (0.01–0.02).

### 3.4 Type-specific predictions are more accurate

In Table 3 we compare the generic *ensnet a* with type-specific *ensnet p*, *ensnet s* and *ensnet n* models, each trained on their corresponding `BioDL` training sets. When tested on the corresponding interaction-specific data sets `BioDL_P_TE`, `BioDL_S_TE` and `BioDL_N_TE`, the specific models consistently obtain performances higher than the interaction-generic *ensnet_a*, as one may expect.

**Table 3:** Performance of *ensnet a*, trained on the *generic* `BioDL_A_TR` data set, compared with the *ensnet p*, *s* and *n* models trained on *type-specific* data sets containing Protein, Small molecule or Nucleotide interaction interfaces, and scored performance on the interaction-specific test sets as indicated.

| Model | Test set | ACC | SPEC | F1 | MCC | AP | AUC |
| --- | --- | --- | --- | --- | --- | --- | --- |
| *ensnet_a* | BioDL_P-TE | 0.828 | 0.902 | 0.289 | 0.192 | 0.248 | 0.733 |
| *ensnet_p* |  | 0.840 | 0.909 | 0.339 | 0.249 | 0.302 | **0.755** |
| *ensnet_a* | BioDL_S-TE | 0.937 | 0.967 | 0.339 | 0.306 | 0.289 | 0.826 |
| *ensnet_s* |  | 0.944 | 0.970 | 0.413 | 0.384 | 0.388 | **0.864** |
| *ensnet_a* | BioDL_N-TE | 0.901 | 0.947 | 0.272 | 0.219 | 0.238 | 0.835 |
| *ensnet_n* |  | 0.921 | 0.957 | 0.418 | 0.376 | 0.399 | **0.894** |

Highest AUC per metric per test set indicated in bold ( $p < 1e-6$ ).

### 3.5 Network architecture matters

We compared the performance of all seven of our models (six sepa-rate architectures and an ensemble) trained on `HHC_TR` and the four `BioDL_*_TR`, yielding 35 trained predictors. Each was applied to their corresponding test sets: three for HHC TR-trained models (Homo TE, Hetero TE and combined `HHC_TE`), and two each for the `BioDL_*_TR` (`BioDL_*_TE` and `ZK448_*_TE`). See SI Tables S3, S4, S5, S6 and S7, for `HHC`, `BioDL_A, _P, _S` and `_N`, respectively, for details. The ensemble *ensnet* predictors perform best on all test sets, as we already saw for the `BioDL_P_TR` models on `BioDL_P_TE` in Table 2.

We further compared our *ensnet hhc*, *ensnet a*, *ensnet s* and *ensnet n* predictors with other published and available state-of-the-art sequence-based predictors *on the same data sets*. The published predictors use various methods and architectures including Random Forest, Logistic Regression, Support Vector Machine, and Neural Networks. Table 4 shows comparisons with SeRenDIP and other PPI predictors benchmarked by [38] on the PPI data sets, and `SCRIBER` and DRNApred on the `ZK448` small molecule and nucleotide (DNA/RNA) data sets, respectively. All predictors mentioned here in Table 4 use similar input features, such as protein length, ASA, RSA, PSSM, and secondary structure predicted from sequence.

**Table 4:** Performance comparison of our *ensnet* models and other state-of-the art methods on applicable test sets.

| Model | Test set | ACC | SPEC | F1 | MCC | AP | AUC |
| --- | --- | --- | --- | --- | --- | --- | --- |
| *Protein-protein interaction (PPI):* |  |  |  |  |  |  |  |
| *ensnet_hhc* | Homo_TE | 0.767 | 0.849 | 0.485 | **0.335** | 0.491 | **0.769**‡ |
| SeRenDIP |  |  |  |  | 0.277 |  | 0.724 |
| *ensnet_hhc* | Hetero_TE | 0.849 | 0.916 | 0.197 | 0.114 | 0.155 | **0.661** * |
| SeRenDIP |  |  |  |  | **0.122** |  | 0.636 |
| *ensnet_a* | ZK448_A_TE | 0.785 | 0.870 | **0.385** | **0.254** | **0.357** | **0.729**‡ |
| SCRIBER † |  | n.a. | **0.896** | 0.333 | 0.230 | 0.287 | 0.715 |
| SSWRF † |  | n.a. | 0.891 | 0.287 | 0.178 | 0.256 | 0.687 |
| CRFPPI † |  | n.a. | 0.887 | 0.266 | 0.154 | 0.238 | 0.681 |
| LORIS † |  | n.a. | 0.887 | 0.263 | 0.151 | 0.228 | 0.656 |
| SPRINGS † |  | n.a. | 0.882 | 0.229 | 0.111 | 0.201 | 0.625 |
| PSIVER † |  | n.a. | 0.874 | 0.191 | 0.066 | 0.170 | 0.581 |
| SPRINT † |  | n.a. | 0.873 | 0.183 | 0.057 | 0.167 | 0.570 |
| SPPIDER † |  | n.a. | 0.870 | 0.198 | 0.071 | 0.159 | 0.517 |
| *Protein-small molecule interaction:* |  |  |  |  |  |  |  |
| *ensnet_s* | ZK448_S_TE | **0.899** | **0.945** | **0.419** | **0.364** | **0.409** | **0.849**‡ |
| SCRIBER |  | 0.874 | 0.931 | 0.278 | 0.209 | 0.259 | 0.706 |
| *Protein-DNA/RNA interaction:* |  |  |  |  |  |  |  |
| *ensnet_n* | ZK448_N_TE | **0.871** | **0.927** | **0.469** | **0.396** | **0.460** | **0.823**‡ |
| DRNAPred (DNA) |  | 0.830 | 0.903 | 0.294 | 0.198 | 0.240 | 0.609 |
| DRNAPred (RNA) |  | 0.814 | 0.894 | 0.230 | 0.124 | 0.248 | 0.547 |

Highest scores per metric per test set indicated in bold; confidence for difference in AUC-ROC with runner-up: \* $p < 0.05$ , ‡ $p < 0.005$ . † metrics according to [38].

For comparing *ensnet hhc* with SeRenDIP, we used exactly the same test sets as used by SeRenDIP. For comparing *ensnet a* with the predictors as published in [38], we exactly followed their testing approach: we calculated average of metrics over 10 subsets of randomly selected 50% of `ZK448_A_TE`, and we only considered PPIs as the to be predicted interactions and all other types as non-interacting. For comparing *ensnet s* with SCRIBER, we randomly selected 10 proteins (UniProt IDs in SI Table S12) from `ZK448_S_TE` and calculated the comparison metrics based on the protein–small molecule binding propensities returned by their webserver. For comparing *ensnet n* with DRNApred, we used all proteins in `ZK448_N_TE` (38 proteins) and calculated the comparison metrics based on the nucleotide interaction propensities returned by their webserver. The cutoff points were selected such that the number of false positives (FPs) and false negatives (FNs) are equal; this affects ACC, SPEC, F1 and MCC. As can be seen from Table 4, our *ensnet hhc*, *ensnet a*, *ensnet s*, and *ensnet n* predictors perform better than the corresponding state-of-the art methods, on virtually all considered metrics.

### 3.6 Feature importance

Figure 4 shows the top 15 ranking of the importance of the features, as measured by SHAP, for one of our models (*ann p*), estimated for 5000 randomly selected amino acids from `ZK448_P`. A SHAP value indicates the contribution of each feature to a residue’s interface prediction; the width of the distribution of a feature’s SHAP values shows its relative importance across the sampled residues. For the aggregated features we show the sum of SHAP values, e.g. `WM_PC` is the sum of SHAP values of `3_wm_PC`, `5_wm_PC`, `7_wm_PC`, and `9_wm_PC`.

**Figure 4:**
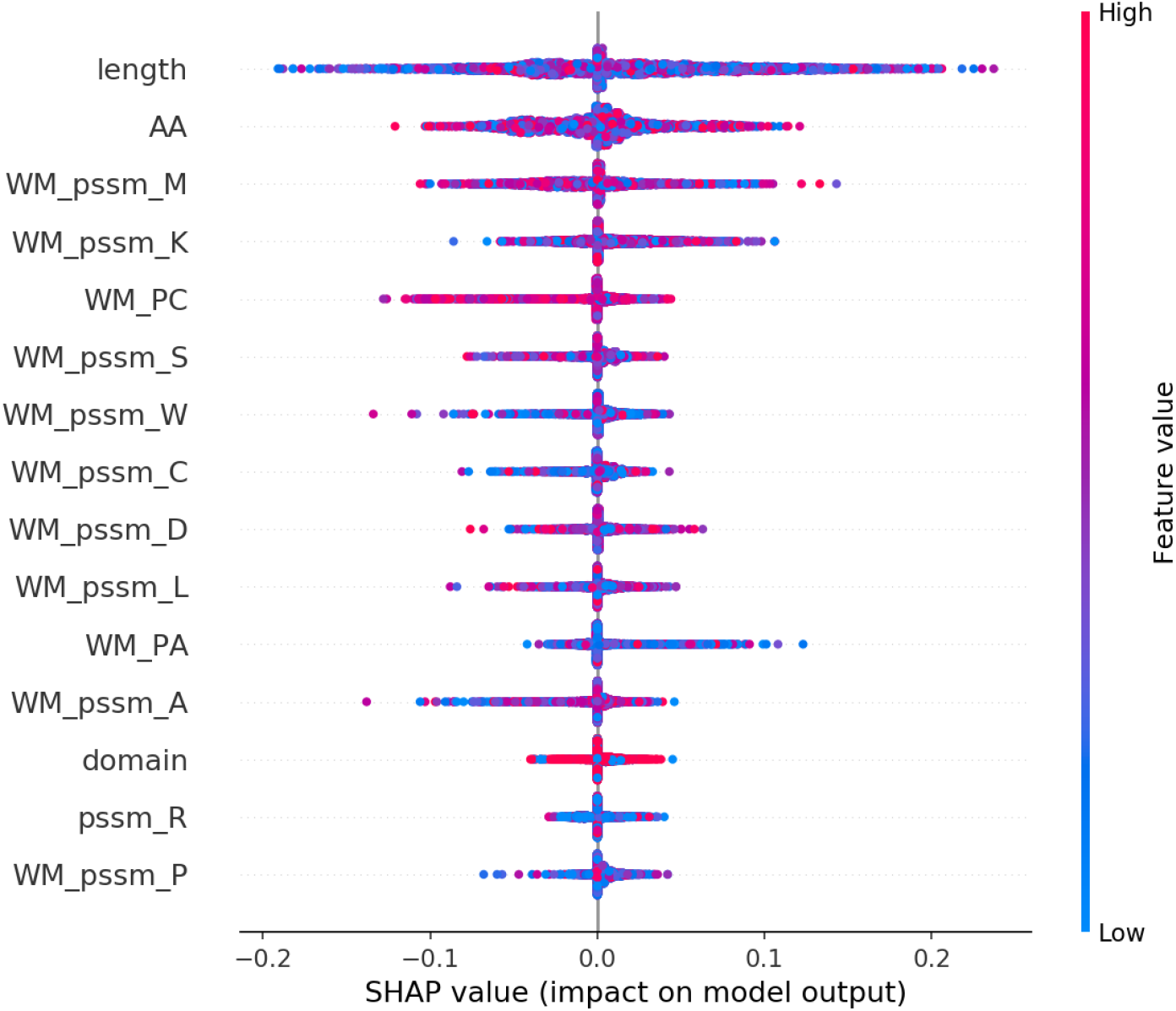
The top 15 ranking of the importance of the features based on the SHAP values for 5000 randomly selected amino acids in `ZK448_P`. Colors represent the input values of a feature: blue for low and red for high values.

The figure also shows the relation between the input value of a feature and its SHAP value. High values of `length` (red dots; residues of longer proteins), in general have lower SHAP values, i.e. lower likelihood of a residue being predicted to be part of an interface. High `WM_PC` values (probability of coil across the windows) also have noteworthy impact on the predictions. SI Fig. S12 shows the correlation of SHAP values for the three secondary structure features: `WM_PA, WM_PB` and `WM_PC`.

## 4 Discussion and conclusion

This work presents an in-depth and systematic comparison of multiple Deep Learning architectures for predicting protein interface residues from sequence, and a series of neural nets, PIPENN, whose ensemble method performs well on generic and type-specific interface prediction tasks, including PPI, small molecule and nucleotide (DNA/RNA) interfaces.

We explored multiple combinations of DL architecture building blocks, such as spatial forms, encoding schemes, network initializations, loss and activation functions, and regularization mechanisms. Selected combinations resulted in six models and an ensemble, which we trained on existing and newly constructed training data sets. Performance was benchmarked on several independent test sets, facilitating fair comparison. Comparing the performance of our models to that of other published and available state-of-the-art sequence-based predictors on the same test sets, shows that our ensemble predictors obtain most accurate predictions on all interface types.

It is worth noting that the different prediction tasks are not equally difficult. We reproduce an earlier observation that homomeric PPI interface residues can be better predicted than heteromeric interfaces, as noted in [9, 37]. Moreover, all architectures predict protein–small molecule and protein–nucleotide interface residues more accurately, while the prediction of protein–protein interface residues appears most difficult. This might be explained by differences in the size, specificity and structural heterogeneity of interfaces involved with these respective types of interaction: nucleotide interfaces are generally much smaller than PPI interfaces, affecting their variety and relative sequence locality; small molecule interactions typically require very specific chemical properties, leading to equally specific amino acids forming their interface; and the structural similarity between two random strands of DNA is much larger than that between two random proteins, so it stands to reason the similarity between (and consequently, predictability of) two protein–DNA interfaces is also greater than the similarity between two protein–protein interfaces.

In summary, we contribute the following: (i) `BioDL`, a protein interaction data set with residue-level and type-specific interface annotation of sufficient size to perform deep learning; (ii) systematic characterization of different combinations of architectural building blocks, and their impact on the predictive performance of resulting neural nets; (iii) the PIPENN suite of neural net models, whose ensemble method outperforms state-of-the-art sequence-based models when it comes to predicting various types of protein interface. This conclusively demonstrates Deep Learning can contribute much to current efforts in computational protein interface prediction. We provide a public repository containing source code, data sets and pre-trained models: https://github.com/ibivu/pipenn/.

## Supporting information

PIPENN supplemental figures and tables

## Acknowledgements

We kindly acknowledge sponsoring by the VU HPC Council and the IT for Research (ITvO) BAZIS Linux computational cluster at the VU University Amsterdam. The authors declare that there is no conflict of interest.

## Author contributions

RH, BS, KAF and SA designed the experiments. HdF, KAF, BS and RH collected the data sets. RH implemented the methods and performed the experiments. RH, BS and KAF analysed and interpreted the results. RH, BS, SA, JH and KAF wrote the article text. All authors revised the article and approved the final version for publication.

