## Supplementary material for "PIPENN: Protein Interface Prediction with an Ensemble of Neural Nets": PIPENN supplemental figures and tables

Bas Stringer [1,#], Hans de Ferrante [1], Sanne Abeln [1], Jaap Heringa [1], K. Anton Feenstra [1], Reza Haydarlou [1,#,*]

[1]IBIVU – Center for Integrative Bioinformatics, Dept. of Computer Science, Vrije Universiteit, Amsterdam 1081HV, The Netherlands

### these authors have contributed equally.

September 3, 2021

#### 1 Data set features and statistcis

- SI Table S1: Train data set statistics
- SI Table S2: Test data set statistics
- SI Table S13: Overview of all 128 features used, and the labels by which they are referenced
- SI Figure S13: The Venn diagram of the training data sets showing common proteins among different protein binding data sets
- SI Figure S14: The Venn diagram of the testing data sets showing common proteins among different protein binding data sets

#### 2 Deep learning architectures

- SI Figure S2: The architecture of the fully connected feed forward neural network (ann)
- SI Figure S3: The architecture of the dilated convolutional neural network (dnet)
- SI Figure S4: The architecture of the U-shape convolutional neural network (unet)
- SI Figure S5: The architecture of the residual neural network (rnet)

- SI Figure S6: The architecture of the recurrent neural network (rnn)
- SI Figure S7: The architecture of the residual recurrent neural network combined with the recurrent neural network (cnet)
- SI Figure S8: The architecture of the ensemble neural network (ensnet)

#### 3 Performance of the DL-architectures

- SI Table S3: Performance of all architectures trained on `HHC_TR` and tested on `Homo_TE`, `Hetero_TE` and `HHC_TE` PPI datasets
- SI Table S4: Performance of all architectures trained on `BioDL_A_TR` and tested on `BioDL_A_TE` or `ZK448_A_TE`, containing annotations for all types of bindings
- SI Table S5: Performance of all architectures trained on `BioDL_P_TR` and tested on `BioDL_P_TE` and `ZK448_P_TE`, containing annotations for PPI
- SI Table S6: Performance of all architectures trained on `BioDL_S_TR` and tested on `BioDL_S_TE` and `ZK448_S_TE`, containing annotations for protein–small molecule bindings
- SI Table S7: Performance of all architectures trained on `BioDL_N_TR` and tested on `BioDL_N_TE` and `ZK448_N_TE`, containing annotations for protein–DNA/RNA bindings
- SI Table S8: Performance of *ensnet_a* trained on validation set of `BioDL_A_TR` and tested on the specified test data sets
- SI Table S9: Performance of *ensnet_p* trained on validation set of `BioDL_P_TR` and tested on the specified test data sets
- SI Table S10: Performance of *ensnet_hhc* trained on validation set of `HHC_TR` and tested on the specified test data sets
- SI Table S12: Ten randomly selected proteins for comparison with SCRIBER
- SI Figure S9: ROC plots of all architectures trained on `BioDL_P_TR` and tested on `ZK448_P_TE`
- SI Figure S11: P/R plots of all architectures trained on `BioDL_P_TR` and tested on `ZK448_P_TE`
- SI Figure S10: Scatter plot of MCCs for all proteins of `BioDL_P_TE`
- SI Figure S12: Distribution of SHAP-values showing the importance of different features w.r.t. the input-values of the feature

#### 4 Run-time statistics

- SI Table S11: Duration (in HH:MM) of training per architecture per data set

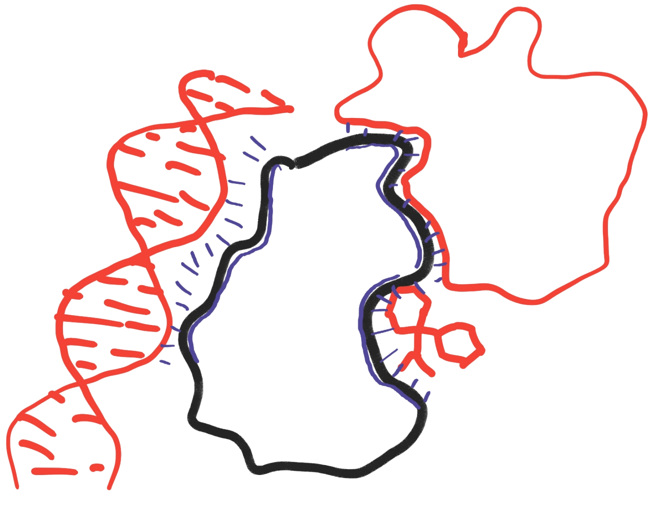

SI Figure S1: Different types of protein binding. A protein (in black) may interact with other molecules (in red) such as nucleic acids (RNA or DNA, left), other proteins (top right), or small molecules (bottom right). Intermolecular interactions are indicated with blue dashes, and the corresponding interface regions on the protein with a blue outline.

SI Table S1: Train data set statistics

| **Training set** | **#Proteins** | **#Residues** | **#Interfaces** |
|---|---|---|---|
| HHC_TR | 407 | 103709 | 20868 (20.1%) |
| BioDL_A_TR | 6832 | 2068793 | 221184 (10.7%) |
| BioDL_P_TR | 4392 | 1279674 | 151670 (11.9%) |
| BioDL_S_TR | 4227 | 1363053 | 67538 (5.0%) |
| BioDL_N_TR | 394 | 131371 | 9552 (7.3%) |

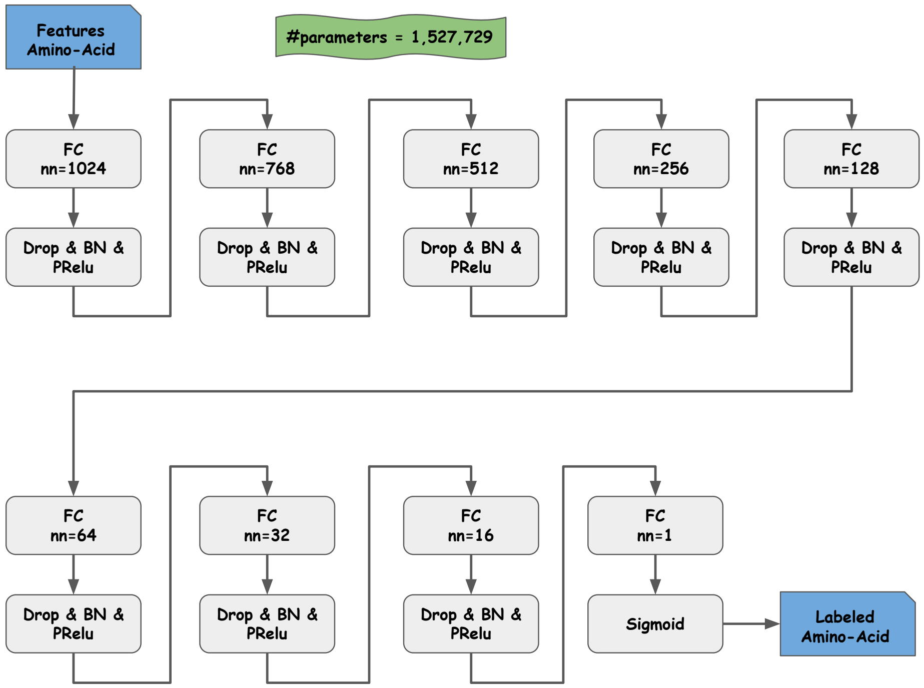


SI Figure S2: The architecture of the fully connected feed forward neural network (ann)

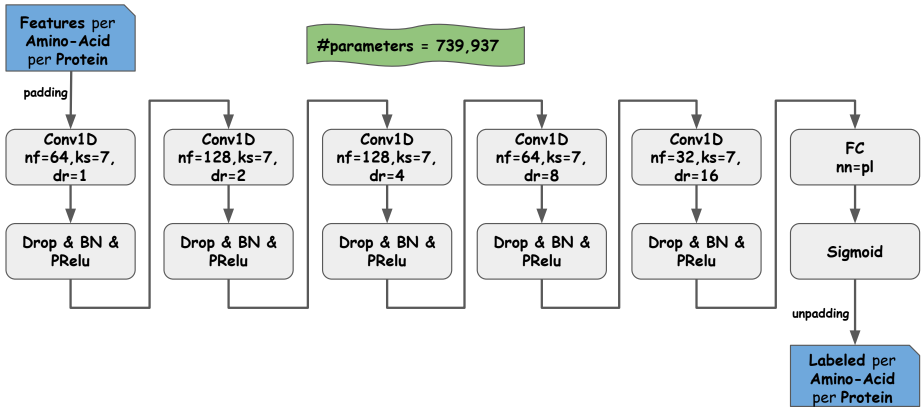


SI Figure S3: The architecture of the dilated convolutional neural network (dnet)

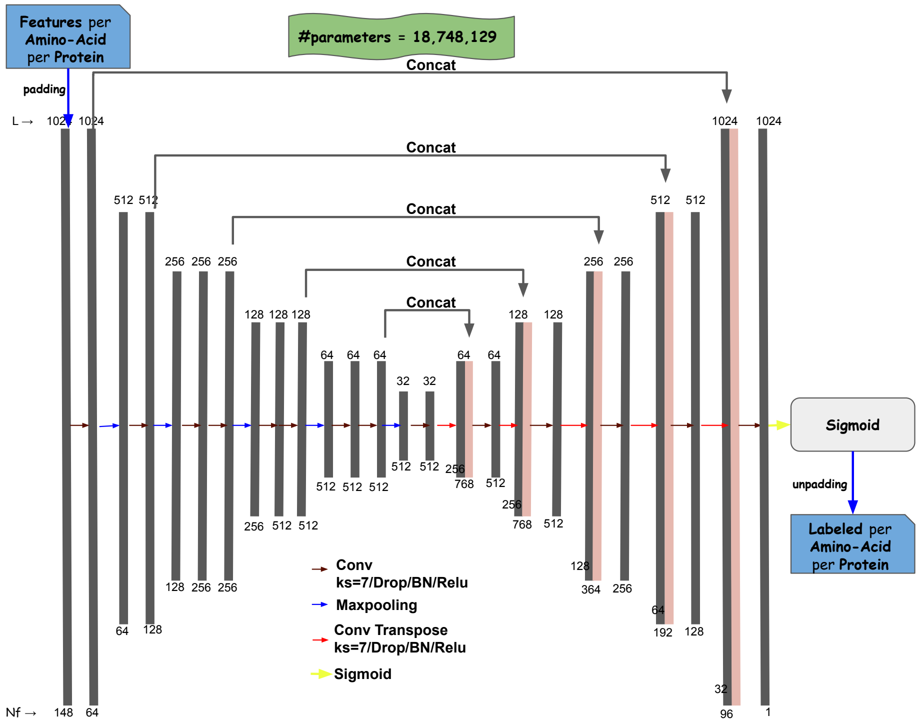


SI Figure S4: The architecture of the U-shape convolutional neural network (unet)

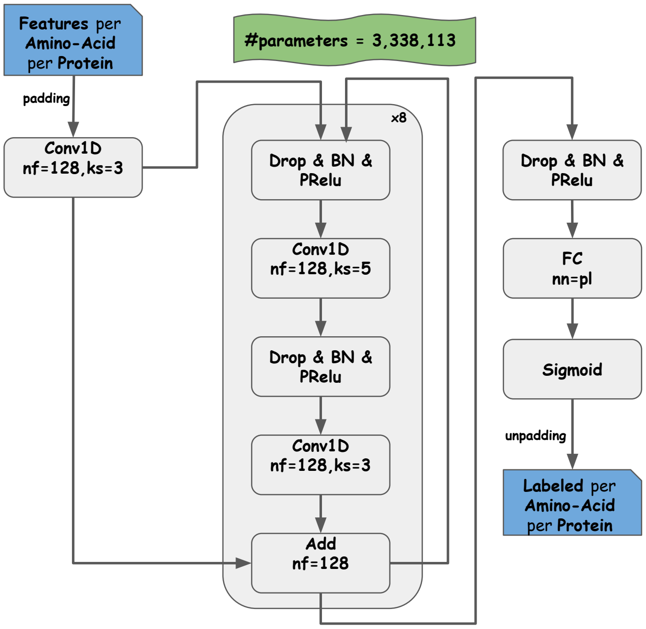


SI Figure S5: The architecture of the residual neural network (rnet)

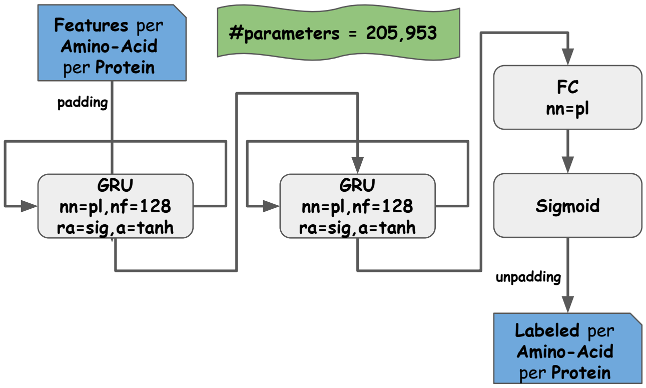


SI Figure S6: The architecture of the recurrent neural network (rnn)

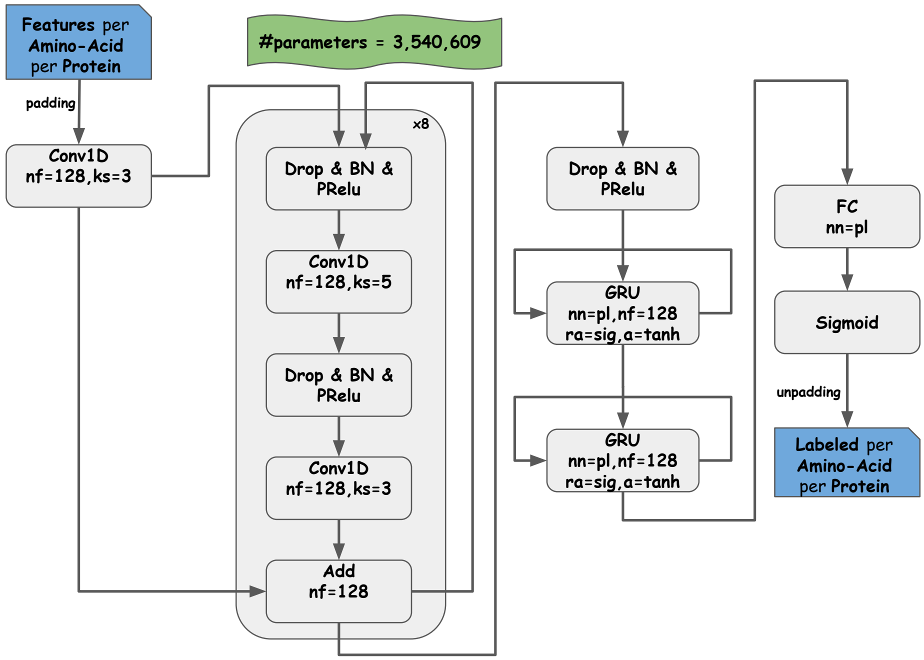


SI Figure S7: The architecture of the residual recurrent neural network combined with the recurrent neural network (cnet)

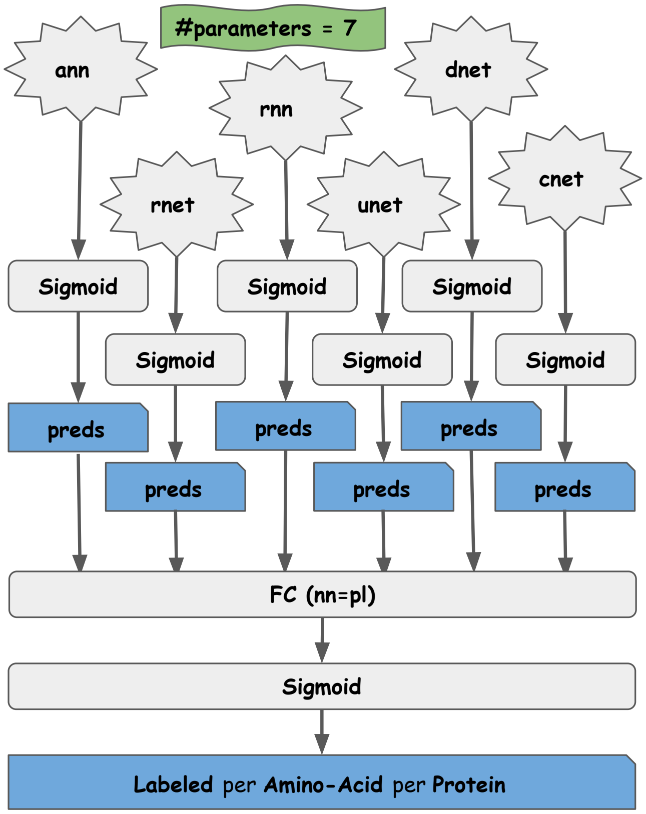


SI Figure S8: The architecture of the ensemble neural network (ensnet)

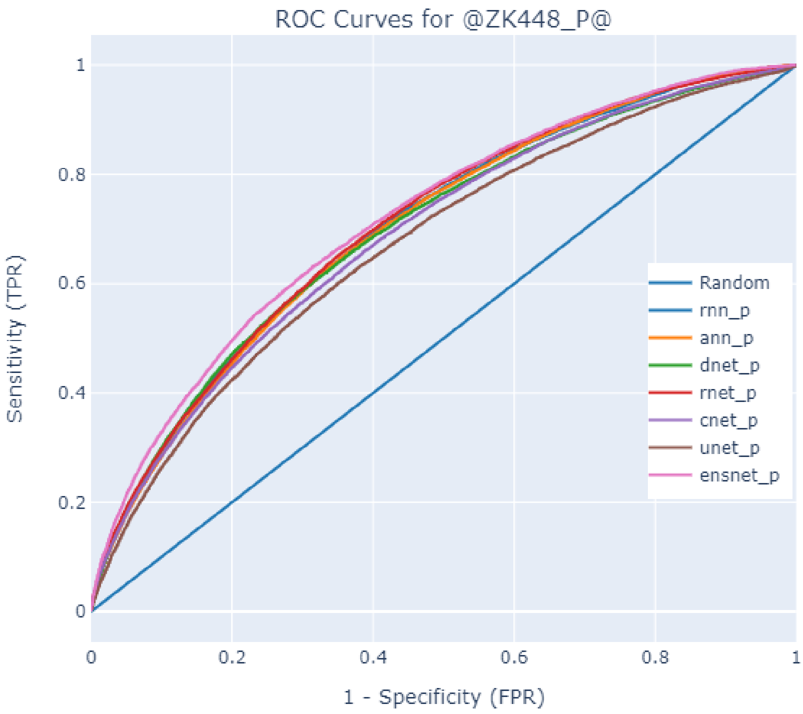


SI Figure S9: ROC plots of all architectures trained on `BioDL_P_TR` and tested on `ZK448_P_TE`

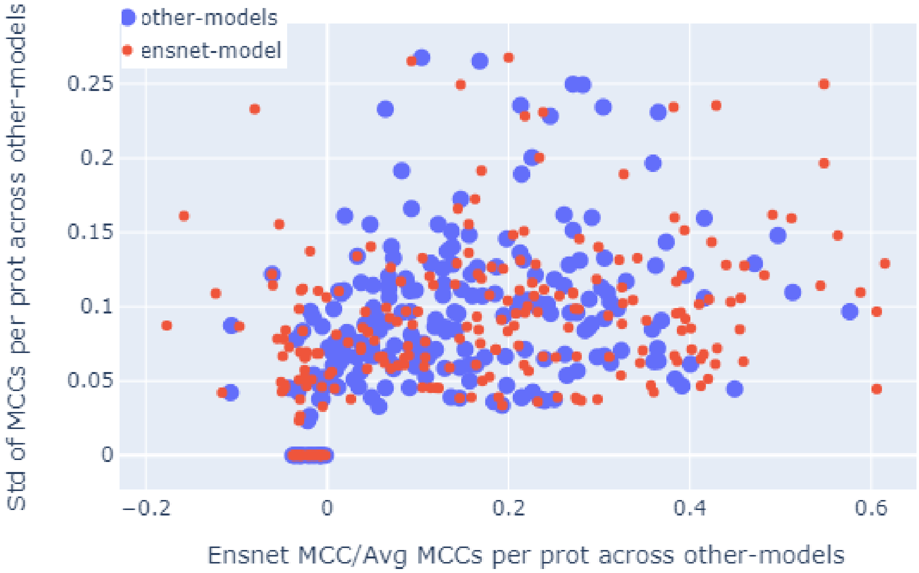


SI Figure S10: Scatter plot of MCCs for all proteins of `BioDL_P_TE` The x-axis of each red/blue dot is the MCC score/average MCC score for a protein, calculated based on the PPI predictions by *ensnet_p*/all other predictors, and the y-axis is the standard deviation of the MCC scores of all other predictors. We can observe that, (1) the MCC scores are the same for all models including *ensnet_p* (zero std. dev, at bottom left), (2) the MCC scores has been distributed among all predictors including *ensnet_p* (y-axis), and (3) the difference of MCC scores between *ensnet_p* and all the other predictors (distance between each blue and red dot on a horizontal line).

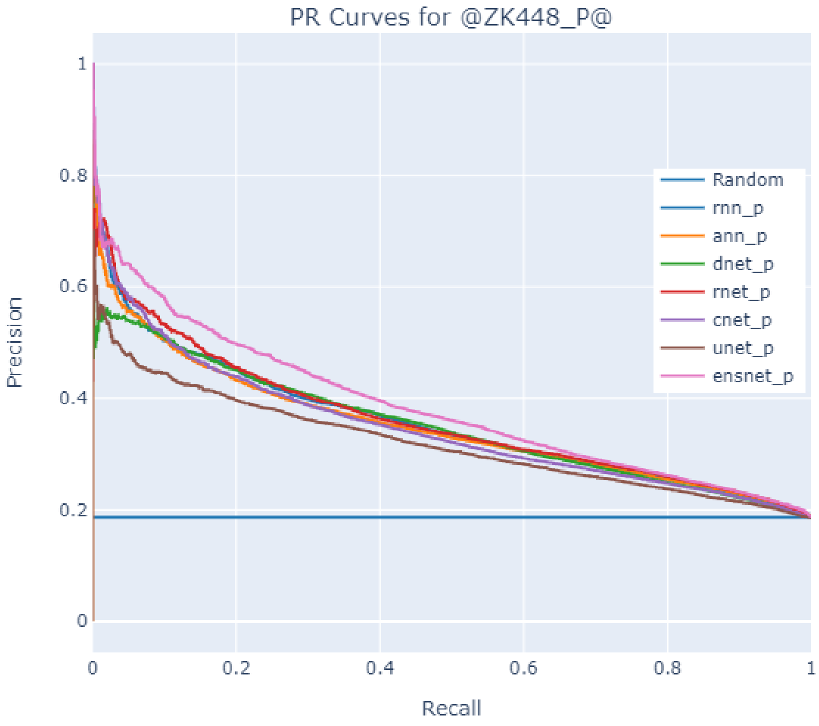


SI Figure S11: P/R plots of all architectures trained on `BioDL_P_TR` and tested on `ZK448_P_TE`

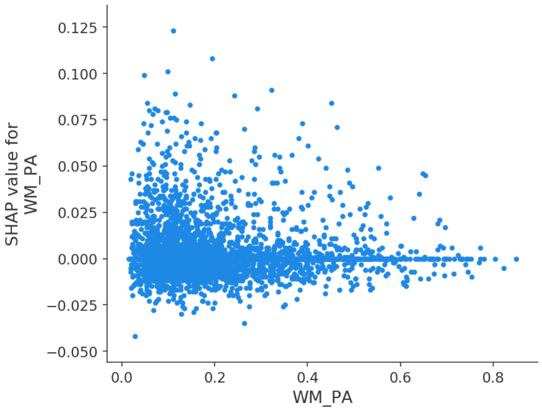


(A)

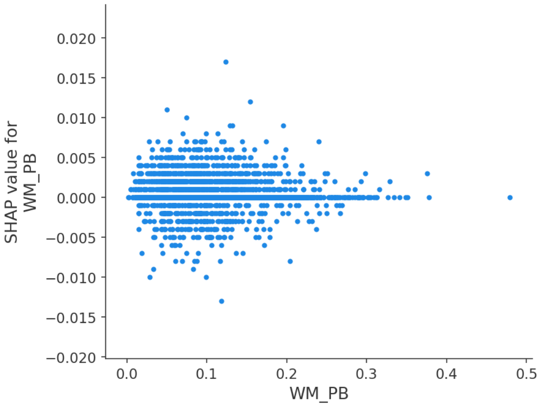


(B)

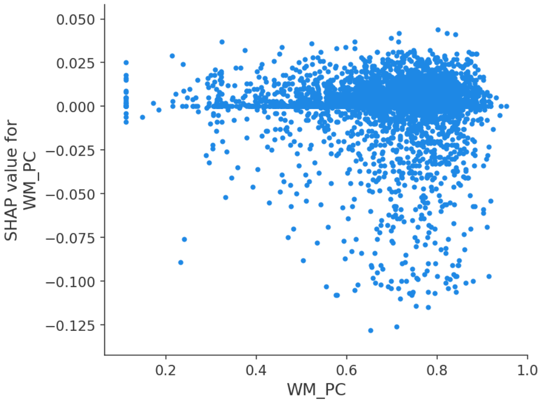


(C)

SI Figure S12: Distribution of SHAP-values showing the importance of different features w.r.t. the input-values of the feature (A) helix. (B) sheet. (C) coil.

SI Table S2: Test data set statistics

| Test set | #Proteins | #Residues | #Interfaces |
|---|---|---|---|
| Homo_TE | 94 | 24488 | 5541 (22.6%) |
| Hetero_TE | 45 | 12991 | 1219 (9.4%) |
| HHC_TE | 139 | 37479 | 6760 (18.0%) |
| BioDL_A_TE | 343 | 100412 | 10951 (10.9%) |
| BioDL_P_TE | 228 | 65150 | 7845 (12.0%) |
| BioDL_S_TE | 202 | 64609 | 3055 (4.7%) |
| BioDL_N_TE | 23 | 7551 | 511 (6.8%) |
| ZK448_A_TE | 448 | 116500 | 22643 (19.4%) |
| ZK448_P_TE | 336 | 84941 | 15810 (18.6%) |
| ZK448_S_TE | 354 | 99318 | 7175 (7.2%) |
| ZK448_N_TE | 38 | 10212 | 1233 (12.1%) |

SI Table S3: Performance of all architectures trained on HHC_TR and tested on Homo_TE, Hetero_TE and HHC_TE PPI datasets

| Architecture | Test set | ACC | SPEC | F1 | MCC | AP | AUC |
|---|---|---|---|---|---|---|---|
| *ensnet_hhc* | Homo_TE | 0.767 | 0.849 | 0.485 | 0.335 | 0.491 | **0.769** |
| *dnet_hhc* | Homo_TE | 0.766 | 0.848 | 0.482 | 0.331 | 0.482 | 0.758 |
| *unet_hhc* | Homo_TE | 0.761 | 0.845 | 0.471 | 0.317 | 0.481 | 0.752 |
| *cnet_hhc* | Homo_TE | 0.754 | 0.841 | 0.458 | 0.299 | 0.436 | 0.732 |
| *rnet_hhc* | Homo_TE | 0.749 | 0.838 | 0.446 | 0.284 | 0.447 | 0.731 |
| *rnn_hhc* | Homo_TE | 0.737 | 0.830 | 0.419 | 0.25 | 0.413 | 0.708 |
| *ann_hhc* | Homo_TE | 0.725 | 0.822 | 0.394 | 0.216 | 0.389 | 0.686 |
| *ensnet_hhc* | Hetero_TE | 0.849 | 0.916 | 0.197 | 0.114 | 0.155 | **0.661** |
| *dnet_hhc* | Hetero_TE | 0.849 | 0.916 | 0.196 | 0.113 | 0.147 | 0.642 |
| *rnet_hhc* | Hetero_TE | 0.841 | 0.912 | 0.154 | 0.066 | 0.132 | 0.640 |
| *rnn_hhc* | Hetero_TE | 0.845 | 0.914 | 0.177 | 0.091 | 0.141 | 0.619 |
| *unet_hhc* | Hetero_TE | 0.840 | 0.912 | 0.152 | 0.064 | 0.133 | 0.614 |
| *ann_hhc* | Hetero_TE | 0.834 | 0.908 | 0.115 | 0.024 | 0.120 | 0.612 |
| *cnet_hhc* | Hetero_TE | 0.841 | 0.912 | 0.154 | 0.066 | 0.127 | 0.604 |
| *ensnet_hhc* | HHC_TE | 0.785 | 0.869 | 0.405 | 0.274 | 0.389 | **0.741** |
| *ann_hhc* | HHC_TE | 0.749 | 0.847 | 0.304 | 0.151 | 0.271 | 0.654 |
| *dnet_hhc* | HHC_TE | 0.784 | 0.868 | 0.403 | 0.272 | 0.381 | 0.733 |
| *unet_hhc* | HHC_TE | 0.779 | 0.865 | 0.388 | 0.253 | 0.365 | 0.719 |
| *rnet_hhc* | HHC_TE | 0.771 | 0.860 | 0.365 | 0.226 | 0.345 | 0.706 |
| *cnet_hhc* | HHC_TE | 0.775 | 0.862 | 0.376 | 0.239 | 0.343 | 0.705 |
| *rnn_hhc* | HHC_TE | 0.766 | 0.857 | 0.352 | 0.21 | 0.332 | 0.688 |

Homo_TE, containing annotations for homomeric PPI; Hetero_TE, containing annotations for heteromeric PPI; HHC_TE, containing annotations for homo- and heteromeric PPI

SI Table S4: Performance of all architectures trained on `BioDL_A_TR` and tested on `BioDL_A_TE` or `ZK448_A_TE`, containing annotations for all types of bindings

| **Architect.** | **Test set** | **ACC** | **SPEC** | **F1** | **MCC** | **AP** | **AUC** |
|---|---|---|---|---|---|---|---|
| *ensnet_a* | `BioDL_A_TE` | 0.853 | 0.917 | 0.327 | 0.245 | 0.296 | **0.762** |
| *rnn_a* | `BioDL_A_TE` | 0.850 | 0.916 | 0.315 | 0.232 | 0.282 | 0.756 |
| *dnet_a* | `BioDL_A_TE` | 0.850 | 0.916 | 0.314 | 0.230 | 0.280 | 0.752 |
| *rnet_a* | `BioDL_A_TE` | 0.850 | 0.916 | 0.313 | 0.229 | 0.276 | 0.745 |
| *cnet_a* | `BioDL_A_TE` | 0.849 | 0.915 | 0.309 | 0.225 | 0.273 | 0.744 |
| *unet_a* | `BioDL_A_TE` | 0.848 | 0.914 | 0.303 | 0.217 | 0.265 | 0.742 |
| *ann_a* | `BioDL_A_TE` | 0.847 | 0.914 | 0.302 | 0.216 | 0.266 | 0.731 |
| *ensnet_a* | `ZK448_A_TE` | 0.775 | 0.860 | 0.422 | 0.283 | 0.414 | **0.736** |
| *rnet_a* | `ZK448_A_TE` | 0.772 | 0.858 | 0.414 | 0.272 | 0.398 | 0.730 |
| *cnet_a* | `ZK448_A_TE` | 0.768 | 0.856 | 0.404 | 0.260 | 0.389 | 0.724 |
| *dnet_a* | `ZK448_A_TE` | 0.768 | 0.856 | 0.403 | 0.260 | 0.391 | 0.723 |
| *rnn_a* | `ZK448_A_TE` | 0.771 | 0.858 | 0.412 | 0.270 | 0.398 | 0.722 |
| *ann_a* | `ZK448_A_TE` | 0.769 | 0.857 | 0.407 | 0.264 | 0.392 | 0.717 |
| *unet_a* | `ZK448_A_TE` | 0.765 | 0.854 | 0.397 | 0.251 | 0.382 | 0.714 |

SI Table S5: Performance of all architectures trained on `BioDL_P_TR` and tested on `BioDL_P_TE` and `ZK448_P_TE`, containing annotations for PPI

| **Architect.** | **Test set** | **ACC** | **SPEC** | **F1** | **MCC** | **AP** | **AUC** |
|---|---|---|---|---|---|---|---|
| *ensnet_p* | `BioDL_P_TE` | 0.840 | 0.909 | 0.339 | 0.249 | 0.302 | **0.755** |
| *dnet_p* | `BioDL_P_TE` | 0.834 | 0.905 | 0.312 | 0.218 | 0.276 | 0.739 |
| *rnn_p* | `BioDL_P_TE` | 0.833 | 0.905 | 0.310 | 0.215 | 0.276 | 0.736 |
| *rnet_p* | `BioDL_P_TE` | 0.833 | 0.905 | 0.309 | 0.215 | 0.279 | 0.735 |
| *cnet_p* | `BioDL_P_TE` | 0.832 | 0.904 | 0.303 | 0.208 | 0.273 | 0.733 |
| *ann_p* | `BioDL_P_TE` | 0.833 | 0.905 | 0.309 | 0.214 | 0.27 | 0.729 |
| *unet_p* | `BioDL_P_TE` | 0.829 | 0.903 | 0.292 | 0.196 | 0.253 | 0.717 |
| *ensnet_p* | `ZK448_P_TE` | 0.775 | 0.862 | 0.397 | 0.259 | 0.384 | **0.718** |
| *rnet_p* | `ZK448_P_TE` | 0.766 | 0.856 | 0.373 | 0.230 | 0.360 | 0.706 |
| *rnn_p* | `ZK448_P_TE` | 0.768 | 0.857 | 0.377 | 0.235 | 0.356 | 0.703 |
| *ann_p* | `ZK448_P_TE` | 0.764 | 0.855 | 0.367 | 0.222 | 0.35 | 0.700 |
| *dnet_p* | `ZK448_P_TE` | 0.768 | 0.857 | 0.378 | 0.235 | 0.350 | 0.697 |
| *cnet_p* | `ZK448_P_TE` | 0.763 | 0.854 | 0.364 | 0.218 | 0.348 | 0.689 |
| *unet_p* | `ZK448_P_TE` | 0.758 | 0.851 | 0.350 | 0.201 | 0.319 | 0.671 |

SI Table S6: Performance of all architectures trained on BioDL_S_TR and tested on BioDL_S_TE and ZK448_S_TE, containing annotations for protein–small molecule bindings

| Architect. | Test set | ACC | SPEC | F1 | MCC | AP | AUC |
|---|---|---|---|---|---|---|---|
| *ensnet_s* | BioDL_S_TE | 0.944 | 0.970 | 0.413 | 0.384 | 0.388 | **0.864** |
| *cnet_s* | BioDL_S_TE | 0.942 | 0.969 | 0.389 | 0.359 | 0.353 | 0.854 |
| *dnet_s* | BioDL_S_TE | 0.942 | 0.969 | 0.395 | 0.365 | 0.356 | 0.852 |
| *rnn_s* | BioDL_S_TE | 0.940 | 0.969 | 0.376 | 0.345 | 0.350 | 0.851 |
| *rnet_s* | BioDL_S_TE | 0.942 | 0.969 | 0.387 | 0.357 | 0.356 | 0.850 |
| *unet_s* | BioDL_S_TE | 0.941 | 0.969 | 0.380 | 0.349 | 0.335 | 0.843 |
| *ann_s* | BioDL_S_TE | 0.938 | 0.967 | 0.352 | 0.320 | 0.291 | 0.830 |
| *ensnet_s* | ZK448_S_TE | 0.916 | 0.955 | 0.424 | 0.380 | 0.402 | **0.842** |
| *cnet_s* | ZK448_S_TE | 0.914 | 0.953 | 0.405 | 0.359 | 0.373 | 0.831 |
| *rnn_s* | ZK448_S_TE | 0.912 | 0.953 | 0.397 | 0.350 | 0.363 | 0.830 |
| *dnet_s* | ZK448_S_TE | 0.913 | 0.953 | 0.403 | 0.356 | 0.371 | 0.827 |
| *rnet_s* | ZK448_S_TE | 0.912 | 0.952 | 0.391 | 0.344 | 0.369 | 0.826 |
| *unet_s* | ZK448_S_TE | 0.913 | 0.953 | 0.403 | 0.356 | 0.368 | 0.822 |
| *ann_s* | ZK448_S_TE | 0.907 | 0.950 | 0.362 | 0.312 | 0.309 | 0.804 |

SI Table S7: Performance of all architectures trained on BioDL_N_TR and tested on BioDL_N_TE and ZK448_N_TE, containing annotations for protein–DNA/RNA bindings

| Architect. | Test set | ACC | SPEC | F1 | MCC | AP | AUC |
|---|---|---|---|---|---|---|---|
| *ensnet_n* | BioDL_N_TE | 0.921 | 0.957 | 0.418 | 0.376 | 0.399 | **0.894** |
| *dnet_n* | BioDL_N_TE | 0.915 | 0.954 | 0.373 | 0.328 | 0.351 | 0.880 |
| *rnet_n* | BioDL_N_TE | 0.915 | 0.954 | 0.375 | 0.33 | 0.339 | 0.879 |
| *rnn_n* | BioDL_N_TE | 0.917 | 0.955 | 0.393 | 0.349 | 0.356 | 0.878 |
| *cnet_n* | BioDL_N_TE | 0.919 | 0.956 | 0.407 | 0.364 | 0.374 | 0.878 |
| *unet_n* | BioDL_N_TE | 0.918 | 0.956 | 0.401 | 0.357 | 0.363 | 0.870 |
| *ann_n* | BioDL_N_TE | 0.913 | 0.953 | 0.358 | 0.311 | 0.323 | 0.865 |
| *ensnet_n* | ZK448_N_TE | 0.871 | 0.927 | 0.469 | 0.396 | 0.460 | **0.823** |
| *ann_n* | ZK448_N_TE | 0.864 | 0.923 | 0.440 | 0.363 | 0.418 | 0.813 |
| *rnet_n* | ZK448_N_TE | 0.865 | 0.923 | 0.443 | 0.367 | 0.408 | 0.813 |
| *rnn_n* | ZK448_N_TE | 0.866 | 0.924 | 0.448 | 0.372 | 0.426 | 0.810 |
| *cnet_n* | ZK448_N_TE | 0.864 | 0.922 | 0.438 | 0.361 | 0.387 | 0.809 |
| *unet_n* | ZK448_N_TE | 0.866 | 0.923 | 0.446 | 0.370 | 0.423 | 0.804 |
| *dnet_n* | ZK448_N_TE | 0.864 | 0.922 | 0.437 | 0.359 | 0.415 | 0.800 |

SI Table S8: Performance of *ensnet_a* trained on validation set of BioDL_A_TR and tested on the specified test data sets

| Test data set | ACC | SPEC | F1 | MCC | AP | AUC |
|---|---|---|---|---|---|---|
| BioDL_A_TE | 0.853 | 0.917 | 0.327 | 0.245 | 0.296 | 0.762 |
| ZK448_A_TE | 0.775 | 0.86 | 0.422 | 0.283 | 0.414 | 0.736 |
| BioDL_P_TE | 0.828 | 0.902 | 0.289 | 0.192 | 0.248 | 0.733 |
| ZK448_P_TE | 0.754 | 0.849 | 0.339 | 0.188 | 0.305 | 0.691 |
| BioDL_S_TE | 0.937 | 0.967 | 0.339 | 0.306 | 0.289 | 0.826 |
| ZK448_S_TE | 0.905 | 0.949 | 0.348 | 0.297 | 0.307 | 0.797 |
| BioDL_N_TE | 0.901 | 0.947 | 0.272 | 0.219 | 0.238 | 0.835 |
| ZK448_N_TE | 0.848 | 0.913 | 0.373 | 0.286 | 0.344 | 0.769 |
| Homo_TE | 0.718 | 0.818 | 0.378 | 0.196 | 0.361 | 0.679 |
| Hetero_TE | 0.855 | 0.92 | 0.229 | 0.149 | 0.186 | 0.692 |

SI Table S9: Performance of *ensnet_p* trained on validation set of BioDL_P_TR and tested on the specified test data sets

| Test data set | ACC | SPEC | F1 | MCC | AP | AUC |
|---|---|---|---|---|---|---|
| BioDL_P_TE | 0.84 | 0.909 | 0.339 | 0.249 | 0.302 | 0.755 |
| ZK448_P_TE | 0.775 | 0.862 | 0.397 | 0.259 | 0.384 | 0.718 |
| Homo_TE | 0.736 | 0.829 | 0.416 | 0.246 | 0.415 | 0.701 |
| Hetero_TE | 0.858 | 0.921 | 0.245 | 0.167 | 0.201 | 0.689 |

SI Table S10: Performance of *ensnet_hhc* trained on validation set of HHC_TR and tested on the specified test data sets

| Test data set | ACC | SPEC | F1 | MCC | AP | AUC |
|---|---|---|---|---|---|---|
| Homo_TE | 0.767 | 0.849 | 0.485 | 0.335 | 0.491 | 0.769 |
| Hetero_TE | 0.849 | 0.916 | 0.197 | 0.114 | 0.155 | 0.661 |
| BioDL_P_TE | 0.798 | 0.885 | 0.163 | 0.049 | 0.152 | 0.597 |
| ZK448_P_TE | 0.761 | 0.853 | 0.358 | 0.212 | 0.332 | 0.685 |

SI Table S11: Duration (in HH:MM) of training per architecture per data set

| Training set | ann | cnet | dnet | ensnet | rnet | rnn | unet |
|---|---|---|---|---|---|---|---|
| HHC_TR | 01:58 | 01:32 | 00:16 | 00:20 | 00:57 | 00:26 | 01:25 |
| BioDL_A_TR | **18:48** | 08:28 | 05:26 | 10:11 | 07:55 | 07:01 | 10:29 |
| BioDL_P_TR | 13:16 | 05:46 | 02:39 | 03:57 | 04:29 | 03:47 | 09:34 |
| BioDL_S_TR | 11:24 | 06:07 | 01:43 | 02:49 | 04:07 | 03:27 | 06:15 |
| BioDL_N_TR | 00:38 | 01:23 | 00:07 | 00:20 | 00:24 | 00:22 | 00:41 |

SI Table S12: Ten randomly selected proteins for comparison with SCRIBER. List of uniprot IDs of 10 proteins that have been randomly selected from ZK448_S_TE (called ZK10_S_TE) and used for comparing the performance of our *ensnet_s* model with the SCRIBER model for protein–small molecule binding task.

| uniprot_id |
|---|
| O43813 |
| Q2GG79 |
| P14174 |
| Q5SHZ3 |
| Q9YGA6 |
| P45568 |
| P13272 |
| A5JTM5 |
| P9WMH3 |
| G9BEX6 |

SI Table S13: Overview of all 128 features used, and the labels by which they are referenced In total, 6 feature types were used: PSSM, RSA, ASA, secondary structure, domain, and the length of query sequence. For local features (PSSM, RSA, ASA, and secondary structure), a mean windowing approach is used, which is indicated by a prefix '`winsize_wm_`', where `winsize` corresponds to (3, 5, 7, 9) of the neighbouring residues.

| Name | Description | Type |
|---|---|---|
| **Global:** | | |
| `length` | the number of amino acids in the protein sequence | integer |
| **Amino Acid type** | | |
| `AA` | the type of an amino acid in the protein sequence | 1-hot |
| **Position Specific Scoring Matrix (PSSM) generated by PSI-BLAST:** | | |
| `PSSM_A...Y` | the profile score for each amino acid | $0 \ldots 1$ |
| **accessible surface area predicted by NetSurfP:** | | |
| `RSA` | predicted relative solvent accessibility | $0 \ldots 1$ |
| `ASA` | predicted absolute solvent accessibility | $0 \ldots 1$ |
| **secondary structure predicted by NetSurfP:** | | |
| `PA` | probability score for $\alpha$-helix | $0 \ldots 1$ |
| `PB` | probability score for $\beta$-sheet | $0 \ldots 1$ |
| `PC` | probability score for coil | $0 \ldots 1$ |
| **Pfam Domain Classification:** | | |
| `domain` | whether or not a residue belongs to a protein domain from Pfam database | 0/1 |

| Feature prefix | `3_wm_` | `5_wm_` | `7_wm_` | `9_wm_` |
|---|---|---|---|---|
| Aggregated over | $-1 \cdots +1$ | $-2 \cdots +2$ | $-3 \cdots +3$ | $-4 \cdots +4$ |

Amino acid can be any of ['A', 'C', 'E', 'D', 'G', 'F', 'I', 'H', 'K', 'M', 'L', 'N', 'Q', 'P', 'S', 'R', 'T', 'W', 'V', 'Y']

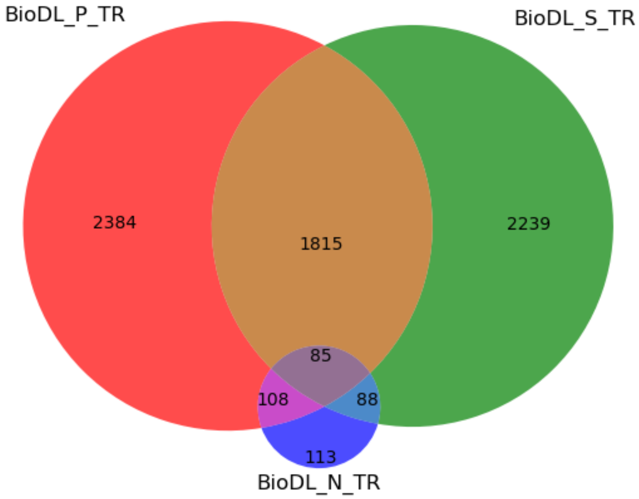


SI Figure S13: The Venn diagram of the training data sets showing common proteins among different protein binding data sets Overlap between **P**rotein, **N**ucleotide and **S**mall molecule interaction annotations in the `BioDL` training set, respectively `BioDL_P_TR`, `BioDL_N_TR` and `BioDL_S_TR`.

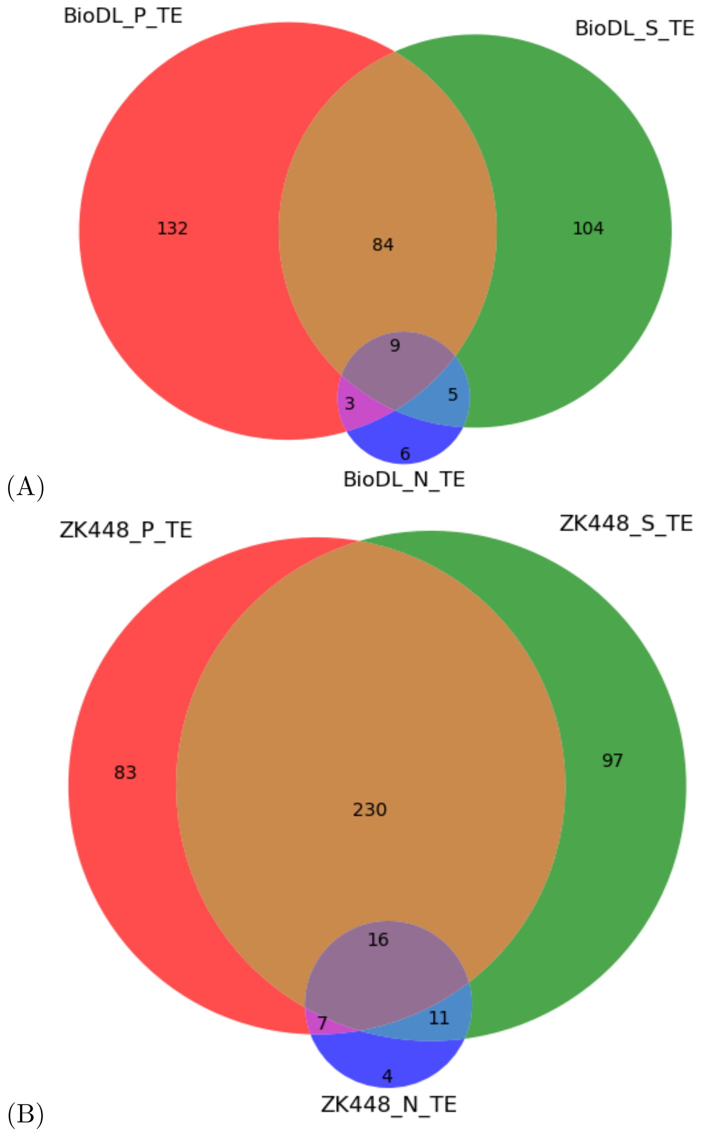


SI Figure S14: The Venn diagram of the testing data sets showing common proteins among different protein binding data sets (A) `BioDL_TE`. (B) `ZK448_TE`.
